# Mutational dynamics and transmission properties of SARS-CoV-2 superspreading events in Austria

**DOI:** 10.1101/2020.07.15.204339

**Authors:** Alexandra Popa, Jakob-Wendelin Genger, Michael Nicholson, Thomas Penz, Daniela Schmid, Stephan W. Aberle, Benedikt Agerer, Alexander Lercher, Lukas Endler, Henrique Colaço, Mark Smyth, Michael Schuster, Miguel Grau, Francisco Martinez, Oriol Pich, Wegene Borena, Erich Pawelka, Zsofia Keszei, Martin Senekowitsch, Jan Laine, Judith H. Aberle, Monika Redlberger-Fritz, Mario Karolyi, Alexander Zoufaly, Sabine Maritschnik, Martin Borkovec, Peter Hufnagl, Manfred Nairz, Günter Weiss, Michael T. Wolfinger, Dorothee von Laer, Giulio Superti-Furga, Nuria Lopez-Bigas, Elisabeth Puchhammer-Stöckl, Franz Allerberger, Franziska Michor, Christoph Bock, Andreas Bergthaler

## Abstract

Superspreading events shape the COVID-19 pandemic. Here we provide a national-scale analysis of SARS-CoV-2 outbreaks in Austria, a country that played a major role for virus transmission across Europe and beyond. Capitalizing on a national epidemiological surveillance system, we performed deep whole-genome sequencing of virus isolates from 576 samples to cover major Austrian SARS-CoV-2 clusters. Our data chart a map of early viral spreading in Europe, including the path from low-frequency mutations to fixation. Detailed epidemiological surveys enabled us to calculate the effective SARS-CoV-2 population bottlenecks during transmission and unveil time-resolved intra-patient viral quasispecies dynamics. This study demonstrates the power of integrating deep viral genome sequencing and epidemiological data to better understand how SARS-CoV-2 spreads through populations.

**Graphical Abstract:** 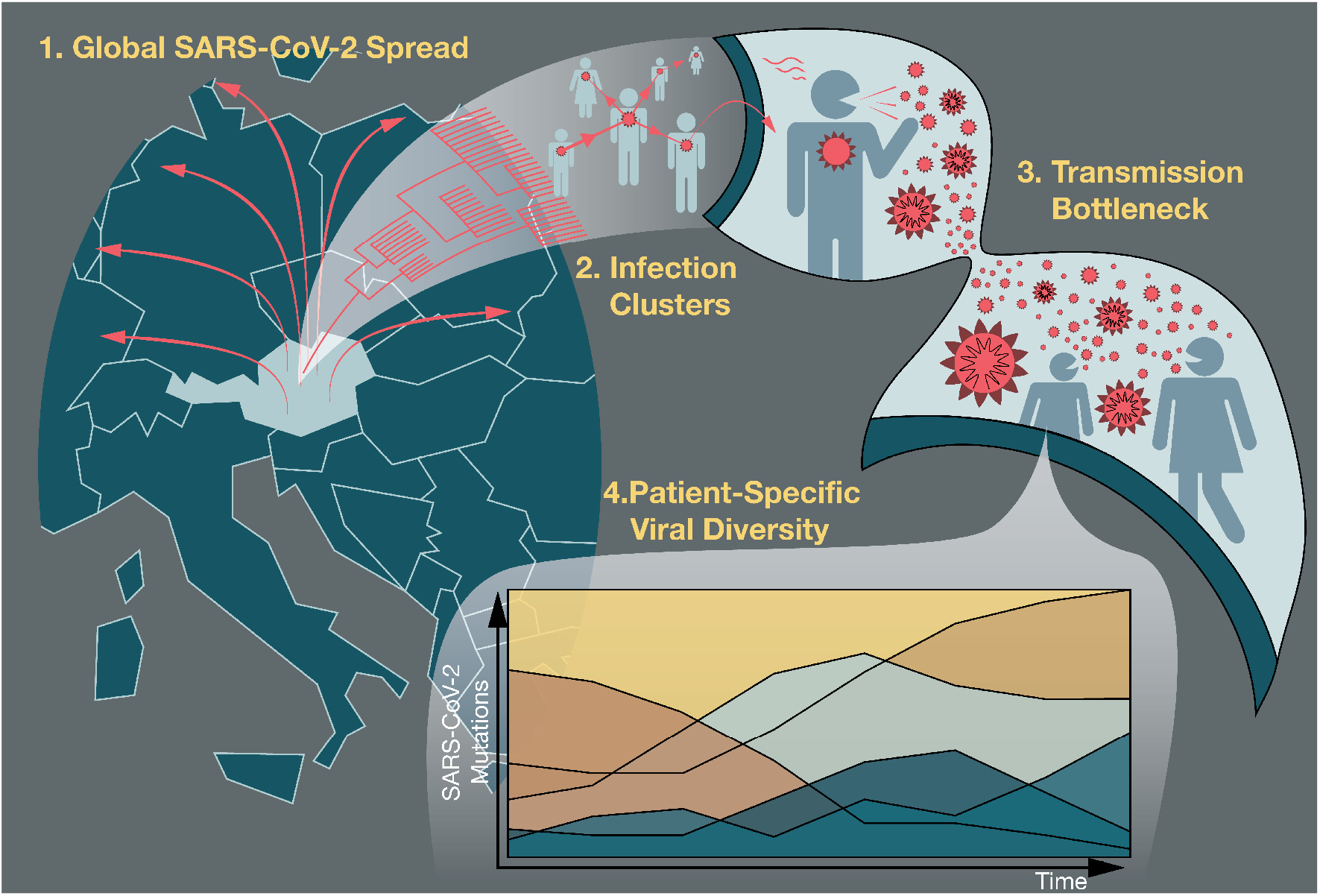

## Introduction

The SARS-CoV-2 pandemic has already infected more than 13 million people in 188 countries, causing 570,375 global deaths as of July 13^th^ 2020 and extraordinary disruptions to daily life and economies (*1*, *2*). The international research community rapidly started to establish diagnostic tools, assess immunological and pathological responses and define risk factors for COVID-19 (*3*–*6*). Clustered outbreaks and superspreading events of SARS-CoV-2 pose a particular challenge to pandemic control (*7*–*10*). However, we still know comparatively little about the fundamental underlying properties of SARS-CoV-2 genome evolution and transmission dynamics within the human population.

During its sweep across the globe, the 29.9 kb-long SARS-CoV-2 genome has accumulated mutations at a rate 2-3 fold lower than those for the SARS, MERS and influenza A viruses (*11*). Acquired fixed mutations enable phylogenetic analyses and have already led to important insights into the origins and routes of SARS-CoV-2 spread (*12*–*15*). Conversely, low frequency mutations and their changes over time within individual patients offer deep insights into the dynamics of intra-host evolution. The resulting viral quasispecies represent diverse groups of variants with different frequencies that form structured populations and maintain high genetic diversity, contributing to fundamental properties of infection and pathogenesis (*16*, *17*).

With a population of 8.8 million people, Austria is located in the center of Europe and operates a highly developed health care system that utilizes a national epidemiological surveillance program. As of June 15^th^ 2020, contact tracing was performed for all 17,082 reported cases that tested positive for SARS-CoV-2, and 6,287 cases could be linked to 502 epidemiological clusters (*18*) (Methods). Due to its prominent role in international winter tourism, Austria has been implicated as a superspreading transmission hub across the European continent. Tourism-associated spread of SARS-CoV-2 from Austria may be responsible for up to half of all imported cases in Denmark, Norway and a considerable share of cases in many other countries including Iceland and Germany (*12*, *19*, *20*).

In this study we phylogenetically and epidemiologically reconstructed major SARS-CoV-2 infection clusters in Austria and analyzed their role in transcontinental virus spread. Moreover, we combined our deep viral genome sequencing data with epidemiologically identified chains of transmissions and family clusters together with biomathematical analyses to study genetic bottlenecks and the dynamics of genome evolution of SARS-CoV-2. Our results provide fully integrated genetic and epidemiological evidence for continental spread of SARS-CoV-2 from Austria and establish fundamental transmission properties of the SARS-CoV-2 in the human population.

## Results

### Phylogenetic-epidemiological reconstruction of SARS-CoV-2 infection clusters in Austria

We sequenced 576 SARS-CoV-2 RNA samples from cases originating from different geographical locations across Austria. The main focus was on the Austrian provinces of Tyrol and Vienna (**Fig. 1A**), given that these two regions led to numerous positive cases (**Fig. S1A**) (*18*). The samples presented in this study capture the first phase of the outbreak (mid-February to mid-March 2020) as well as the peak of infections in Austria (**Fig. 1A**), and cover a balanced sampling across multiple epidemiological and clinical parameters including age, sex and viral load (**Fig. S1B-C**). Samples from both swabs (nasal, oropharyngeal) and secretions (tracheal, bronchial) were included, in order to investigate not only the evolutionary dynamics within the population, but also within individuals (**Fig. S1D**). We assembled SARS-CoV-2 genome sequences, constructed phylogenies and identified low frequency mutations based on high-quality sequencing results with >5 million reads per sample and >80% of mapped viral reads **(Fig. S2A-B**). Our pipeline was validated by experimental controls involving sample titration and technical sample replicates (**Fig. S2C-E**).

**Fig. 1.**
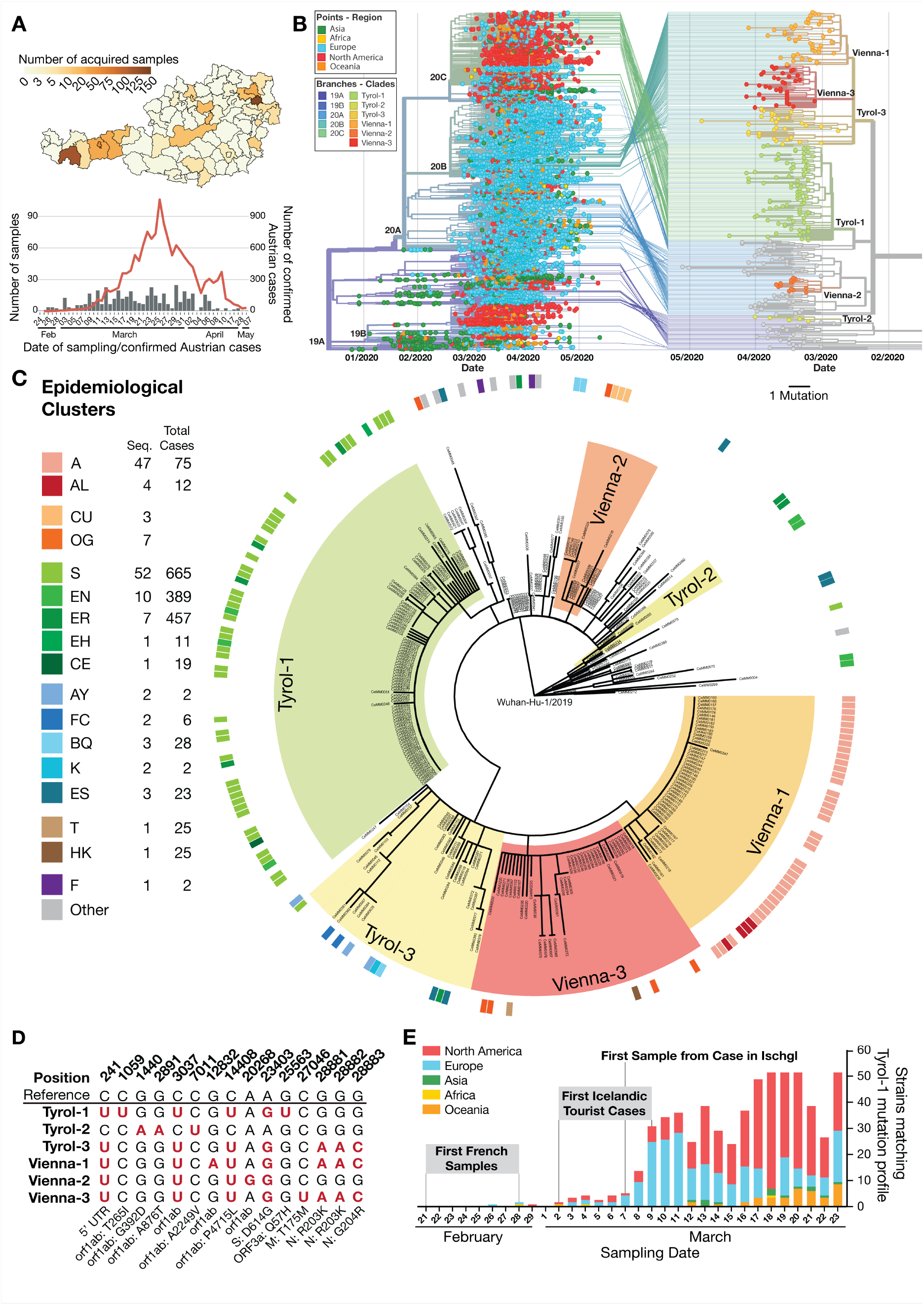
Phylogenetic-epidemiological reconstruction of SARS-CoV-2 infection clusters in Austria. (**A**) Number of acquired samples per district in Austria (top) and sampling dates of samples that underwent viral genome sequencing in this study (bottom), plotted in context of all confirmed cases (red line) in Austria (bottom). (**B**) Connection of Austrian strains to global clades of SARS-CoV-2. Points indicate the regional origin of a strain in the time-resolved phylogenetic tree from 7,695 randomly subsampled sequences obtained from GISAID including 305 Austrian strains sequenced in this study (left). Lines from the global phylogenetic tree (left) to the time-resolved phylogenetic tree of all Austrian strains obtained in this study (right) indicate the phylogenetic relation and Nextstrain clade assignment of Austrian strains. Color schemes of branches represent assignment to clades defined by Nextstrain (left) or phylogenetic clusters of Austrian strains (right). (**C**) Phylogenetic tree of SARS-CoV-2 strains from Austrian COVID-19 patients sequenced in this study. Phylogenetic clusters were identified based on characteristic mutation profiles in patients. Cluster names indicate the most abundant location of patients based on epidemiological data. The circular color code indicates the epidemiological cluster assigned to patients based on contact tracing. (**D**) Mutation profiles of phylogenetic clusters identified in this study. Positions with characteristic mutations compared to the reference sequence “Wuhan-Hu-1” (GenBank: MN908947.3) are highlighted in red. Details regarding the affected genes or genomic regions and the respective codon and amino acid change are given below the table. (**E**) Timeline of the emergence of strains matching the mutation profile of the Tyrol-1 cluster in the global phylogenetic analysis by geographical distribution with additional information from European phylogenetic reconstruction.

To investigate the link between local outbreaks in Austria and the global pandemic, we performed phylogenetic analysis of 305 SARS-CoV-2 genomes from the Austrian cases (>96% genome coverage, >80% aligned viral reads) and 7,695 global genomes from the GISAID database (**Fig. 1B, Table S1**). Our analysis revealed six distinct phylogenetic clusters defined by fixed mutation profiles that were mainly present in the Tyrol region (Tyrol-1, Tyrol-2, Tyrol-3) and in Vienna (Vienna-1, Vienna-2, Vienna-3) (**Fig. 1B**). These clusters are related to the global clades 19A, 20A, 20B and 20C (**Fig. S3A**). Our largest phylogenetic cluster, “Tyrol-1” (**Fig. S3B**), whose cases are closely linked to the ski-resort Ischgl, was assigned to clade 20C (**Fig. 1B**). This clade is predominantly populated by strains from North America (**Fig. S3A**).

Integration of the phylogenetic analysis of Austrian SARS-CoV-2 sequences with epidemiological data from contact tracing resulted in strong overlap of these two lines of evidence (**Fig. 1C, Table S2**). All sequenced samples from the epidemiological cluster A mapped to the relatively homogenous phylogenetic cluster Vienna-1 (**Fig. 1C**) with an index patient with reported travel history to Italy. Of note, both clusters Tyrol-1 and Vienna-1 originate from crowded indoor events (i.e. Apré Ski bar; spinning class), by now known origins for superspreading events.

The epidemiological cluster S, which includes resident and travel-associated cases to the ski-resort Ischgl or the related valley Paznaun, largely mapped to the phylogenetic cluster Tyrol-1 (**Fig. 1C**). While different SARS-CoV-2 strains circulated in the region of Tyrol, this data suggests that the epidemiological cluster S originated from a single strain with a characteristic mutation profile leading to a large outbreak in this region (**Fig. 1D**). To elucidate the possible origin of the SARS-CoV-2 strain giving rise to this cluster, we searched for strains matching its mutation profile among global SARS-CoV-2 sequences (**Fig. 1D-E**, **Fig. S3C**). We found that all strains in clade 20C matched the mutation profile of the Tyrol-1 cluster in our phylogenetic analysis (**Fig. 1E**, **Fig. S3C**). To reveal possible transmission lines between European countries at that time, we performed phylogenetic analysis using all 7,675 high-quality European SARS-CoV-2 sequences sampled before March 31 that were available in the database GISAID. Using this approach, we identified several samples matching the Tyrol-1 cluster mutation profile from a local outbreak in the region Hauts-de-France in the last week of February (**Fig. 1E**, **Fig. S3C**)(*21*). Introduction events of this SARS-CoV-2 strain to Iceland by tourists with a travel history to Austria were reported as early as March 2 (**Fig. 1E, S3C**) (*12*), indicating that this strain was already present in Ischgl in the last week of February. These findings suggest that the emergence of the cluster Tyrol-1 coincided with the local outbreak in France (**Fig. 1E**) and with the early stages of the severe outbreak in Northern Italy (*22*). The viral genomes observed in the Tyrol-1 cluster were closely related to those observed among the Icelandic cases with a travel history to Austria (**Fig. S3D-E**) (*12*). *Vice versa*, many of the Icelandic strains with a “Tyrol-1” mutation profile had reported an Austrian or Icelandic exposure history (**Fig. S3F**). Together, these observations and epidemiological evidence support the notion that the SARS-CoV-2 outbreak in Austria propagated to Iceland. Moreover, the emergence of these strains coincided with the emergence of the global clade 20C (**Fig. S3C**). One week after the occurrence of SARS-CoV-2 strains with this mutation profile in France and Ischgl, an increasing number of related strains based on the same mutation profile could be found across continents (**Fig. 1E**) where they established new local outbreaks, for example in New York City (*13*). As a popular international destination attracting thousands of tourists from European countries and overseas, Ischgl may have played a critical role as transmission hub for the spread of clade 20C strains to Europe and North America (**Fig. S3G-H**).

### Dynamics of low frequency and fixed mutations in clusters

Next, we sought to gain insights into the fundamental processes of SARS-CoV-2 infection by integrative analysis of viral genomes. Mutational analysis of SARS-CoV-2 genomes from Austrian cases revealed that more than half of the observed substitutions were non-synonymous, with the most frequent non-synonymous mutations occurring in nsp6, ORF3a and ORF8 (**Fig. S4A-B**). An analysis of the mutational signatures in the 7,695 global strains and the Austrian subset of SARS-CoV-2 isolates showed a non-homogeneous mutational pattern dominated by C>U, G>U and G>A substitutions (**Fig. S4C**). Moreover, we observed increased mutation rates in the 3’ and 5’ UTRs (**Fig. S4D**). We found that 30% of the positions in the genome (9,194 total positions) harbored variants among the sequenced strains from Austria and we identified mutational hotspots for both high and low-frequency mutations (**Fig. 2A, Fig. S5A-B**). Among these, 8,842 positions exhibit only low frequency mutations (<0.50), while four positions (241, 3037, 14408, 23403) demonstrate fixation of the mutation in more than 50% of samples. We also identified 33 positions with alternative alleles being fixed in more than three samples and exhibiting a frequency <0.50 in at least two other samples (**Fig. 2A**). We confirmed the non-homogeneous mutational pattern of fixed mutations in our low-frequency data, suggesting that the same biological and evolutionary forces are at work for both types of mutations (**Fig. S5C-E**).

**Fig. 2:**
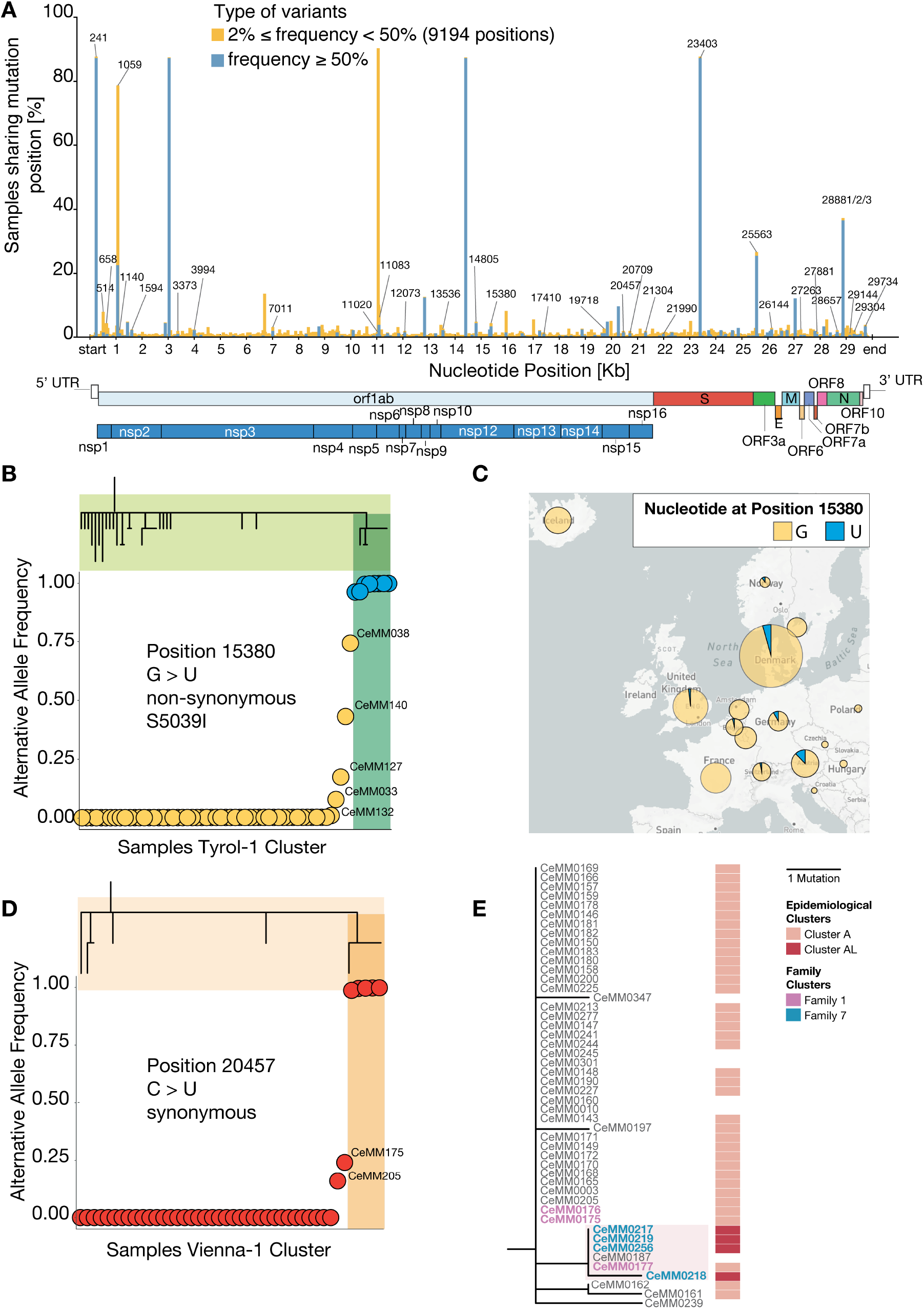
Dynamics of low frequency and fixed mutations in superspreading clusters. Percentage of samples sharing detected (≥ 0.02) mutations across genomic positions. For each of the 9,194 positions harbouring an alternative allele, the percentage of samples with high (≥0.50) or low (≥0.02 and ≤0.50) frequency are reported in dark blue and orange, respectively. (**B**) Allele frequency of non-synonymous mutation G > U, at position 15,380 across samples in the phylogenetic cluster Tyrol-1. This variant has been observed both as low frequency variant and as fixed mutation, the latter defining a phylogenetic subcluster (dark green). (**C**) Proportion of European samples with a reference (yellow) or alternative (blue) allele at position 15,380. (**D**) Allele frequency of synonymous mutation C > U, at position 20,457 across samples of the Vienna-1 phylogenetic cluster. This variant is fixed and defines a phylogenetic subcluster (dark orange) as part of the broader Vienna-1 cluster. (**E**) Phylogenetic tree (genetic divergence) of patient samples in the cluster Vienna-1, overlaid with epidemiological clusters and family-related information.

Based on our phylogenetic analysis, we found a sub-cluster inside the phylogenetic Tyrol-1 cluster, defined by a fixed non-synonymous G>U mutation at position 15,380 (**Fig. 2B**). This mutation is absent from most of the other Austrian cases but was detected at low and intermediary frequencies in other cases of the Tyrol-1 cluster. Interestingly, around the emergence of this mutation, sequences sharing the same mutational profile (i.e. Tyrol-1 haplotype and G>U at position 15,380) appeared in other European countries including Denmark and Germany (**Fig. 2C**). Similarly, a synonymous fixed C>U mutation at position 20,457 defines a subcluster inside the phylogenetic Vienna-1 cluster (**Fig. 2D**). Potential functional effects of this mutation are unknown although it is predicted to slightly alter the locally stable RNA region with a destabilized central fold (**Fig. S6C-D**). The cases from this subcluster intersect with members of two families (families 1 and 7) (**Fig. 2E**, see also **Fig. 3A**). Three members of family 1 tested positive for SARS-CoV-2 on March 8^th^ and were epidemiologically assigned to cluster A. Yet, their viral sequences exhibit a wide range of C>U mutation frequencies at position 20,457 (0.00, 0.24, and 1.00 respectively) (**Fig. 2D-E**). Conversely, four members of family 7, who tested positive for SARS-CoV-2 between March 16^th^ and 22^nd^, were epidemiologically assigned to cluster AL and also show a fixed U nucleotide at position 20,457 (**Fig. 2D-E**). This data indicates the emergence of a fixed mutation within a family, and provides phylogenetic evidence to link previously disconnected epidemiological clusters. Together, these results from two superspreading events (Tyrol-1, Vienna-1) demonstrate the power of deep viral genome sequencing in combination with detailed epidemiological data for observing viral mutation on their way from emergence at low frequency to fixation.

**Fig. 3:**
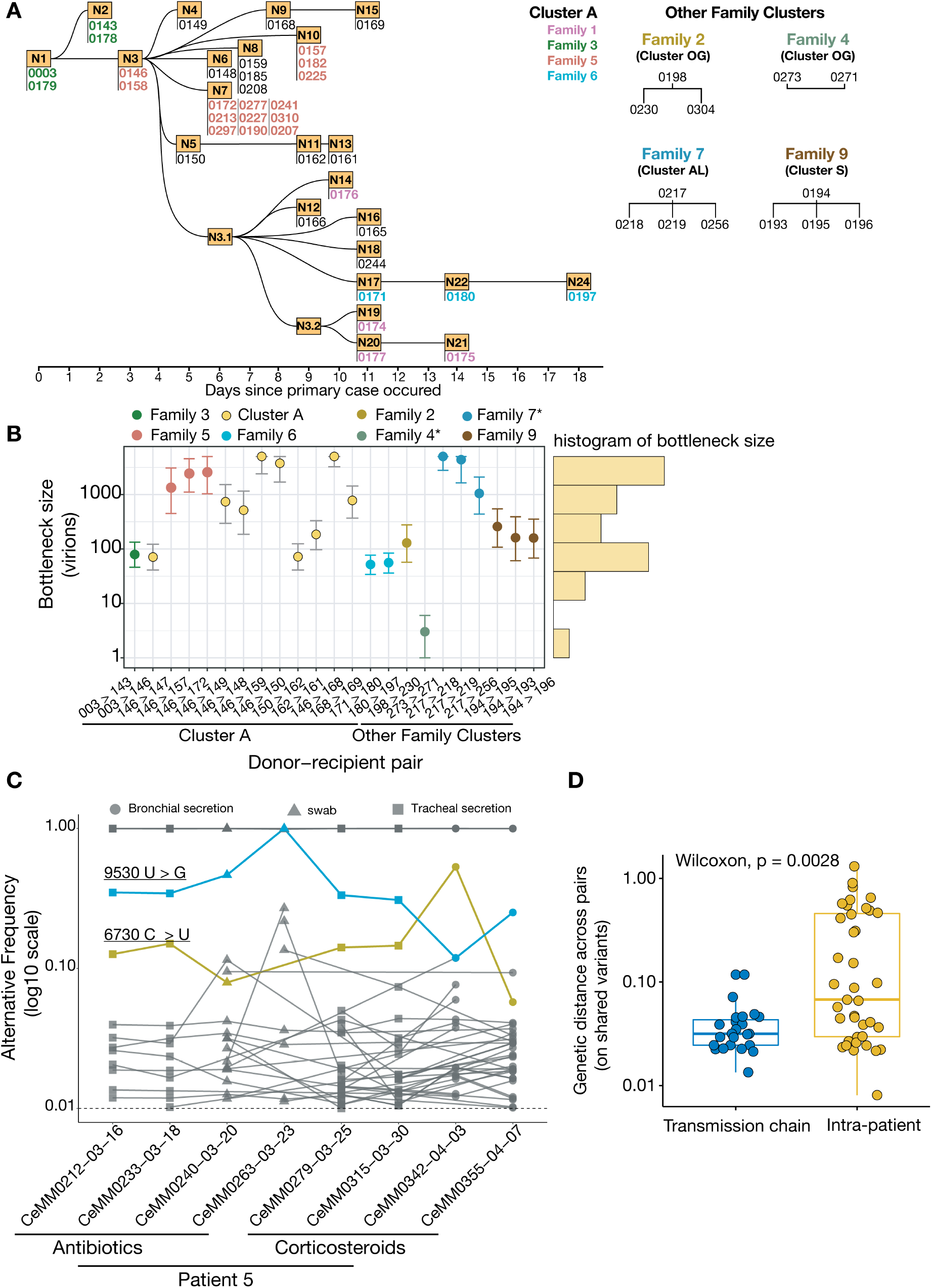
Impact of transmission bottlenecks and intra-host evolution on SARS-CoV-2 mutational dynamics. (**A**) Schematics of the time-related patient interactions across the epidemiological cluster A. Each node represents a case and links between the nodes are epidemiologically confirmed direct transmissions. Samples sequenced from the same individual are reported under the corresponding node. Cases corresponding to the same family are color-coded accordingly. Additional families, unrelated to cluster A, and their epidemiological transmission details are also reported. (**B**) Bottleneck size (i.e. number of virions that initiate the infection in a recipient) estimation across donor-recipient pairs based on **Fig. 3A** and ordered according to the timeline of cluster A for the respective pairs. For patients with multiple samples, the earliest sample was considered for bottleneck size inference. Centered dots are maximum likelihood estimates, with 95% confidence intervals. A * for families 4 and 7 indicates that the infection order was inferred as detailed in the Methods. (**C**) Alternative allele frequency of mutations across available time points for patient 5. Only variants with frequencies >0.01 and shared between at least between two time points are shown. Two mutations increasing in frequency are color coded. (**D**) Genetic distance values of mutation frequencies between donor-recipient pairs (**Fig. 3A-B**) (transmission chains, n=23) and intra-patient consecutive time points (n=39) (**Fig. 3C**, **Fig. S7B**). Only variants seen in two samples are considered.

### Impact of transmission bottlenecks and intra-host evolution on SARS-CoV-2 mutational dynamics

The emergence and potential fixation of mutations in the viral quasispecies depend on interhost bottlenecks and intra-host evolutionary dynamics (*23*, *24*). To test the individual contributions of these forces, we first investigated the transmission dynamics between known pairs of donors and recipients by inferring the number of virions initiating the infection, also known as the genetic bottleneck size (*23*, *25*). Our set of SARS-CoV-2 positive-tested cases comprised twenty-three epidemiologically confirmed donor-recipient pairs (**Fig. 3A**, **Fig. S7A**). Using a beta-binomial method to quantify the bottleneck size (*25*), we found a median bottleneck size of 514 virions (ranging from 3 to greater than 5000) (**Fig. 3B**). The observed relatively large bottleneck size of SARS-CoV-2 is driven by the stability of mutation frequencies across transmissions (**Fig. S7A**).

Next, we investigated the dynamics of intra-host evolution by using time-resolved viral sequences from nine longitudinally sampled patients. These patients were subject to different medical treatments and four of them succumbed due to COVID-19 related complications (**Table S3**). We observed diverse mutation patterns across individual patients and time (**Fig. 3C, Fig. S7B**). Samples from patients 2, 3, and 10 showed a small number of stable low-frequency mutations (≥ 0.02 and ≤0.50), while patients 1, 4, 5, 7, 8 and 9 exhibited higher variability including the fixation of individual mutations (**Fig. 3C, Fig. S7B**). The patient-specific dynamics of viral mutation frequencies may indicate influences of both host-intrinsic factors such as immune responses and overall state of their health as well as extrinsic factors such as different treatment protocols. Finally, we examined the genetic distance between samples obtained across donor - recipient pairs and serially acquired patient samples. This analysis revealed that genetic divergence was greater between the consecutive samples within individual patients than during inter-host transmission (**Fig. 3D**), suggesting that viral intrahost evolution has a potentially strong impact on the landscape of fixed mutations.

## Discussion

Unprecedented global research efforts are underway to match the rapid pace of the COVID-19 pandemic around the globe and its pervasive impact on health and socioeconomics. These efforts include the genetic characterization of SARS-CoV-2 to track viral spread and to dissect the viral genome as it undergoes changes in the human population. Here, we leveraged deep viral genome sequencing in combination with national-scale epidemiological workup to reconstruct Austrian SARS-CoV-2 clusters that played a substantial role in the international spread of the virus. Notably, our time-resolved study shows how emerging low-frequency mutations of SARS-CoV-2 become fixed in local clusters with subsequent spread across countries, thus connecting viral mutational dynamics within individuals and across populations. Exploiting epidemiologically well-defined clusters and families, we were able to determine the inter-human genetic bottleneck for SARS-CoV-2, i.e. the number of virions that start the infection and produce progeny in the viral population, at around 10^1^-10^3^. Our quantified bottlenecks are based on a substantial number of defined donor-recipient pairs and in agreement with recent studies implying larger bottleneck sizes for SARS-CoV-2 compared to estimates for influenza A virus (*23*, *26*–*29*). These bottleneck sizes correlate inversely with higher mutation rates of influenza virus as compared to SARS-CoV-2. Of note, estimates of viral bottleneck size are likely influenced by many parameters including virus-specific differences and stochastic evolutionary processes (*29*). Successful viral transmission also depends on other factors including the rate of decay of viral particles, availability of susceptible cells, the host immune response and co-morbidities (*23*, *30*). To better understand the mechanisms at work, future investigations will need to probe these factors in the context of viral intra-host diversity across body compartments and time (*31*)(*32*) and assess how the underlying viral population structures act together and influence genome evolution of SARS-CoV-2 (*33*, *34*).

This study underscores the value of tightly integrated epidemiological and molecular sequencing approaches to provide the high spatiotemporal resolution necessary for public health experts to track pathogen spread effectively. This enables the retrospective identification of SARS-CoV-2 chains of transmissions and international hotspots such as the phylogenetic cluster Tyrol-1 (*15*, *35*–*37*). Our data also show that all but cluster Tyrol-2 carry the prevalent D614G mutation in the S protein (*38*), supporting the notion of multiple introduction events to Tyrol. Moreover, our phylogenetic analysis of cluster Vienna-1 allowed us to uncover previously unknown links between epidemiological clusters. This observation was subsequently confirmed by extended contact tracing, demonstrating deep viral genome sequencing directly contributes to public health efforts.

The time has come to implement the technical capacities and interdisciplinary framework for prospective near real-time tracking of SARS-CoV-2 infection clusters by integrating approaches which combine viral phylogenetic and epidemiological evidence as well as possible complementary data such as serological testing (*39*). Such inter-disciplinary platforms will be particularly relevant for the prevention of superspreading events and the assessment of the effectiveness of pandemic containment strategies in order to improve the preparedness for anticipated recurrent outbreaks and resurgences of SARS-CoV-2 as well as future pandemics (*40*).

## Online Materials and Methods

### Sample collection and processing

Patient samples were obtained from the Medical Universities of Vienna Institute of Virology, Medical University of Innsbruck Institute of Virology, Medical University of Innsbruck Department of Internal Medicine II, Central Institute for Medical-Chemical Laboratory Diagnostics Innsbruck, Klinikum Wels-Grieskirchen and the Austrian Agency for Health and Food Safety (AGES). Samples were obtained from individuals with a suspected SARS-CoV-2 infection, individuals that were tested positive and followed up during the course of disease or individuals that are either related and/or had close contact to a person that had previously tested positive. Sample types include oropharyngeal swabs, nasopharyngeal swabs, tracheal secretion, bronchial secretion, serum, plasma and cell culture supernatants. RNA was extracted using the following commercially available kits by adhering to the manufacturer’s instructions: MagMax (Thermo Fischer), EasyMag (bioMérieux), AltoStar Purification Kit 1.5 (Altona Diagnostics), MagNA Pure LC 2.0 (Roche), MagNA Pure Compact (Roche) and QIAsymphony (Qiagen). Viral RNA was reverse transcribed with Superscript IV Reverse Transcriptase (ThermoFisher, 18090010). The resulting cDNA was used to amplify viral sequences with modified primer pools from the Artic Network Initiative (*41*). PCR reactions were pooled and subjected to Next Generation Sequencing.

### Sample sequencing

Amplicons were cleaned-up with AMPure XP beads (Beckman Coulter) with a 1:1 ratio. Amplicon concentrations were quantified with the Qubit Fluorometric Quantitation system (Life Technologies) and the size distribution was assessed using the 2100 Bioanalyzer system (Agilent). Amplicon concentrations were normalized, and sequencing libraries were prepared using the NEBNext Ultra II DNA Library Prep Kit for Illumina (New England Biolabs) according to manufacturer’s instructions. Library concentrations again were quantified with the Qubit Fluorometric Quantitation system (Life Technologies) and the size distribution was assessed using the 2100 Bioanalyzer system (Agilent). For sequencing, samples were pooled into equimolar amounts. Amplicon libraries were sequenced on NovaSeq 6000 (Illumina) in 250-base-pair, paired-end mode.

### Sequencing data processing and analysis

Following demultiplexing, fastq files containing the raw reads were inspected for quality criteria (base quality, N and GC content, sequence duplication, over-represented sequences) using the FASTQC (v. 0.11.8) (*42*). Trimming of adapter sequences was performed with BBDUK from the BBtools suite (http://jgi.doe.gov/data-and-tools/bbtools). Overlapping read sequences in a pair were corrected for with BBMERGE function from the BBTools.

Read pairs were mapped on the combined Hg38 and SARS-CoV-2 genome (GenBank: MN908947.3) using the BWA-MEM software package (v 0.7.17) (*43*). Only reads mapping uniquely to the SARS-CoV-2 viral genome were extracted. Primer sequences were removed after mapping by masking with iVar (*44*). From the viral reads BAM file, the consensus FASTA file was generated using the Samtools (v 1.9) (*45*), mpileup, Bcftools (v 1.9) (*46*), and SEQTK (https://github.com/lh3/seqtk). For calling low frequency variants the viral read alignment file was realigned using the Viterbi method provided by LoFreq (v 2.1.2) (*47*). After adding InDel qualities low frequency variants were called using LoFreq. Variant filtering was performed with LoFreq and Bcftools (v 1.9) (*48*). Annotations of the variants was performed with SnpEff (v 4.3) (*49*) and SnpSift (v 4.3) (*50*).

### Technical controls

Two synthetic RNA genomes (MT007544.1, #102019, Twist Bioscience; MN908947.3, #102024, Twist Biosciene) were titrated in increasing ratios (0.1%, 1%, 5%, 10%, 90%, 100%) and subjected to cDNA synthesis and PCR amplification as described. These controls are important for assessing the limit of low frequency detection across samples. RNA was extracted and processed independently from sample CeMM0001 to serve as a technical control for PCR processing. Amplicons from the CeMM0008 sample were sequenced twice in order to assess the potential biases introduced by the sequencing step.

### Epidemiological analyses and identification of SARS-CoV-2 infection clusters

In Austria, the Department of Infection Epidemiology & Surveillance at AGES has the legal mandate for contact tracing as part of the epidemiological investigation of the COVID-19 outbreak. Accumulations of cases within a certain time-period in a certain region are called clusters. The aim of epidemiological analysis is to show how an outbreak spreads within the population: To do this, one tries to identify sources of infection and chains of transmission of the cases through personal interviews with diseased or positively tested people (= cases). The aim of contact tracing is to rapidly identify potentially newly infected persons who may have come into contact with existing cases, in order to reduce further onward transmission. The epidemiological SARS-CoV2 clusters were defined as groups of SARS-CoV2 cases, aggregated by time and common exposure. The required information for cluster finding and resolution in chains of transmission was collected during the official case-contact tracing by the public health authorities, resulting in identification of the most likely source cases and successive cases of the index cases. Contact tracing was performed according to technical guidance relating to this measure produced by the European Centre for Disease Prevention and Control (ECDC). [European Centre for Disease Prevention and Control (ECDC). Contact tracing: public health management of persons, including healthcare workers, having had contact with COVID-19 cases in the European Union – second update 2020 [8 April 2020]. (https://www.ecdc.europa.eu/sites/default/files/documents/Contact-tracing-Public-health-management-persons-including-healthcare-workers-having-had-contact-with-COVID-19-cases-in-the-European-Union%E2%80%93second-update_0.pdf]).

### Phylogenetic analysis and inference of transmission lines

Phylogenetic analysis was conducted using the Augur package (version 7.0.2) (*51*). We compiled a randomly subsampled dataset of 7695 full length viral genomes with high coverage (<1% Ns) that were available from GISAID (https://www.gisaid.org/, 2^nd^ of June) and 305 sequences obtained in this publication. Sequences from GISAID were filtered for entries from human hosts with complete sampling dates. Metadata information for patient age and sex were excluded from the analysis. Multiple sequence alignments (MSA) were performed using mafft (*52*). A masking scheme for homoplasic and highly ambiguous sites was applied to avoid bias in the following phylogenetic analysis as discussed elsewhere [N. De Maio, C. Walker, R. Borges, L. Weilguny, G. Slodkowicz, and N. Goldman, “Issues with SARS-CoV-2 sequencing data,” *virological.org*.]. We reconstructed the phylogeny with the augur pipeline using IQ-Tree (*52*) and further processed the resulting trees with treetime to infer ancestral traits of the nodes (*53*). Phylogenetic trees were rooted with the genome of “Wuhan-Hu-1/2019”. The same workflow was repeated for phylogenetic reconstruction of all high-quality European strains before March 31^st^ 2020 available in the GISAID database by June 7^th^ 2020 (7675). Clade annotations for global trees were adapted from nextstrain.org (https://github.com/nextstrain/ncov/blob/master/config/clades.tsv), clusters of Austrian strains were identified based on shared mutation profiles and patient location from epidemiological data.

### Mutational profiles

Inter-host mutations were reconstructed using the augur pipeline to infer nucleotide changes at the internal nodes (*51*). Positions reported as highly homoplasic were masked, including the first 55 and the last 100 nucleotides [N. De Maio, C. Walker, R. Borges, L. Weilguny, G. Slodkowicz, and N. Goldman, “Issues with SARS-CoV-2 sequencing data,” *virological.org*.]. The consequence type of the mutations was annotated using a customized implementation of the Ensembl Variant Effect Predictor (VEP version 92) using the first SARS-CoV-2 sequenced genome (NCBI ID: NC_045512v2) as a reference. The mutational profile was obtained as the normalized count of the number of mutations in each of the 192 trinucleotide changes. To account for the genomic composition of the SARS-CoV-2 virus we also divided each triplet probability by the total number of available triples in the SARS-CoV-2 reference genome. For the intra-host analysis, the process to obtain the mutational spectra panels was the same as intra-host but using the low frequency variant calling output (3090 mutations across 404 Austrian samples with alleles frequencies between 0.05 and 0.5). The mutational profile was computed following the same rationale as for the inter-host variants.

### Genome-wide mutation rate analysis

We aimed to assess whether the variation in the rate of single nucleotide substitution along the SARS-CoV-2 genome can be solely explained by its tri-nucleotide composition. We devised a statistical test performing local estimations of the deviation from the expectation of the observed number of mutations with respect to the expected based on the tri-nucleotide composition of a particular region of the genome. We first counted the total number of non-protein affecting mutations (i.e., synonymous variants and upstream/downstream gene variants) that has been observed across sequenced viral genomes of infected individuals. The focus on non-protein affecting mutations aims to lessen the potential positive selection bias derived from their effect into the coding parts of the viral genome. We did not consider mutations in masked sites (see filtering of mutations for further information about masked sites). We next assigned to each reference nucleoside a probability of mutation of the three alternatives based on its tri-nucleotide context (5’ and 3’ nucleosides) and the relative probability of mutation derived from the 7,695 samples from GSAID. Then we performed N (N=10^6^) randomizations of the same number of observed mutations distributing them along the SARS-CoV-2 genome according to their mutational probability. Protein-affecting mutations were not randomized, and masked sites were not available to the randomization. We then divided the 29,903 bp of the viral genome into 10 windows of 1kb (except the last window with 903 bps). Analogously, in the zoom-in analysis, we divided the first and last 1kb window of the viral genome into 10 windows of 100 bp. For each window we estimated the mean and standard deviation number of simulated mutations within the window. Finally, for each window we estimated the deviation from the expectation using a log-likelihood test (i.e., G-test goodness of fit), where we compared the observed number of mutations in the window versus the mean simulated number.

### RNA secondary structure prediction

To address the question whether mutations that have been observed in the Austrian SARS-CoV-2 samples have an influence on the RNA structure of the virus we performed computational predictions at the secondary structure level with the ViennaRNA package (*54*). We started with characterizing locally stable RNA structures in the SARS-CoV-2 reference genome NC 045512.2 with RNALfold. We required that the underlying sequences were not longer than 150 nt and we targeted thermodynamic stability by selecting only regions whose free energy z score was at least −3 among 1000 dinucleotide shuffled sequences of the same sequence composition. This approach yielded 386 locally stable RNA secondary structures spread throughout the SARS-CoV-2 genome, which were then intersected with 12 unique fixed mutations found in Austrian samples, i.e. positions 241, 1059, 3037, 12832, 14408, 15380, 20457, 23403, 24642, 25563, 27046, and 28881-28883. This approach resulted in seven hits which were subsequently analyzed in detail. We performed single sequence minimum free energy (MFE) structure predictions for both the reference and the mutation variants. In addition, we assessed for each region the level of structural conservation within a set of phylogenetically related viruses. Here we were particularly interested in finding evidence for covariation in stacked helices. Typical covariation patterns are compensatory mutations, i.e. cases where a mutation in one nucleotide is compensated by a second mutation of its pairing partner, such as a GC base-pair being replaced by an AU pair. Likewise, consistent mutations comprise cases where only one nucleotide is exchanged, thereby maintaining the base-pair, e.g. GC to GU. We characterized orthologous regions in selected Sarbecovirus species with Infernal (*55*), produced structural multiple sequence nucleotide alignments with locARNA (*56*)and computed consensus structures with RNAalifold (*57*). In addition, each block was analyzed for structural conservation by RNAz (*58*).

### Bottleneck estimation

We first set out to estimate the transmission bottleneck sizes for each donor-recipient pair. Our analysis is based on the beta-binomial method presented in Leonard et al (*25*). For a given variant present in the donor, this method assumes that the number of transmitted virions carrying the variant is binomially distributed with the bottleneck size as the number of trials and success probability as the variant frequency in the donor. Following transmission, the viral population during early infection is modeled as a linear birth-death process, implying that the proportion of the viral population descended from any virion in the bottleneck population is beta-distributed. Using this model for the change in variant frequencies between donor and recipient pairs and assuming independence of mutations leads to the likelihood model of Leonard et al. Maximum likelihood analysis then provides the bottleneck statistics. Error bars denote 95% confidence intervals, determined by a likelihood ratio test. This method was applied to variants in the frequency range [0.01,0.95]. Due to the high depth of our study we use the approximate version of the beta-binomial method.

### Infection networks

For those families with a known index patient, this patient was assumed to have infected all others in the family. For families with an unknown index patient, we used the likelihoods generated by the beta-binomial bottleneck estimation method in the following way. For a proposed infection network, we obtained the maximum likelihood bottleneck estimate for each edge (donor-recipient pair), unless a variant with frequency greater than 0.95 is lost, in which case the likelihood was set to 0. The likelihood of this network, maximized over possible bottleneck sizes, is then the product of the likelihood for each edge. Building all infection networks for a given family and determining the likelihood of each network then allows selection of the maximum likelihood network. In other words, we jointly maximized the likelihood of the network and bottleneck sizes (a bottleneck size for each edge). Only star-shaped graphs were considered, i.e. a network in which a single index patient infects all other patients in the family. This method neglects de-novo mutations, which should be incorporated for larger scale networks.

### Genetic Distance

For shared mutations with a defined donor to recipient transmission (**Fig. 3B**), we determined those mutations present in both samples and calculated their absolute difference in frequency. Similarly, we made the same computations between time consecutive pairs for serially sampled patients. If multiple samples were obtained on the same day, the sample with lowest Ct value was considered. Note that the time-consecutive pairs had differing number of days between samples. To these genetic distances obtained from the shared variants we added the sum of the frequencies of the variants detected in only one of the pairs of shared samples; i.e. we calculated the l1-norm of the variant frequencies. Statistical difference between the genetic distances from transmission pairs versus consecutive pairs from serially sampled patients, was determined by a Wilcoxon (one-sided) rank-sum test.

## Acknowledgments

We thank the Biomedical Sequencing Facility at CeMM for assistance with next-generation sequencing. We thank Gernot Walder, Peter Obrist, Rainer Gattringer, Christian Paar, Gregor Hörmann for providing samples and Tobias Pahlke for support with computing cluster. We thank the Tourism office Paznaun – Ischgl for statistical data.

This project received funding from the Vienna Science and Technology Fund (WWTF) as part of the WWTF COVID-19 Rapid Response Funding 2020.

## Author contributions

Study design and manuscript writing (AP, JWG, AB), data analysis (AP, JWG, MN, DS, BA, AL, LE, HC, MSm, MSc, MG, FM, OP, ZK, MS, SM, MB, MTW, GSF, NLB, FA, FM, CB, AB), assay design and experimental sample processing (TP, BA, AL, MSe, JL), sample provision and data collection (SWA, WB, EP, JHA, MRF, MK, AZ, PH, MN, GW, DvL, EPS), supervision (CB, AB), coordination (AB).

## Competing interests

Authors declare no competing interests.

## Data and materials availability

Virus sequences are deposited in the GISAID database (see **Table S1**). All phylogenetic trees used in this study are available for visualization under the following URLs: Global build: https://nextstrain.org/community/bergthalerlab/SARS-CoV-2/NextstrainAustria;

European build with European strains before 31 March: https://nextstrain.org/community/bergthalerlab/SARS-CoV-2/EarlyEurope

Build with Austrian strains used for phylogenetic analysis: https://nextstrain.org/community/bergthalerlab/SARS-CoV-2/OnlyAustrian

## Supplementary Figures

**Fig. S1.**
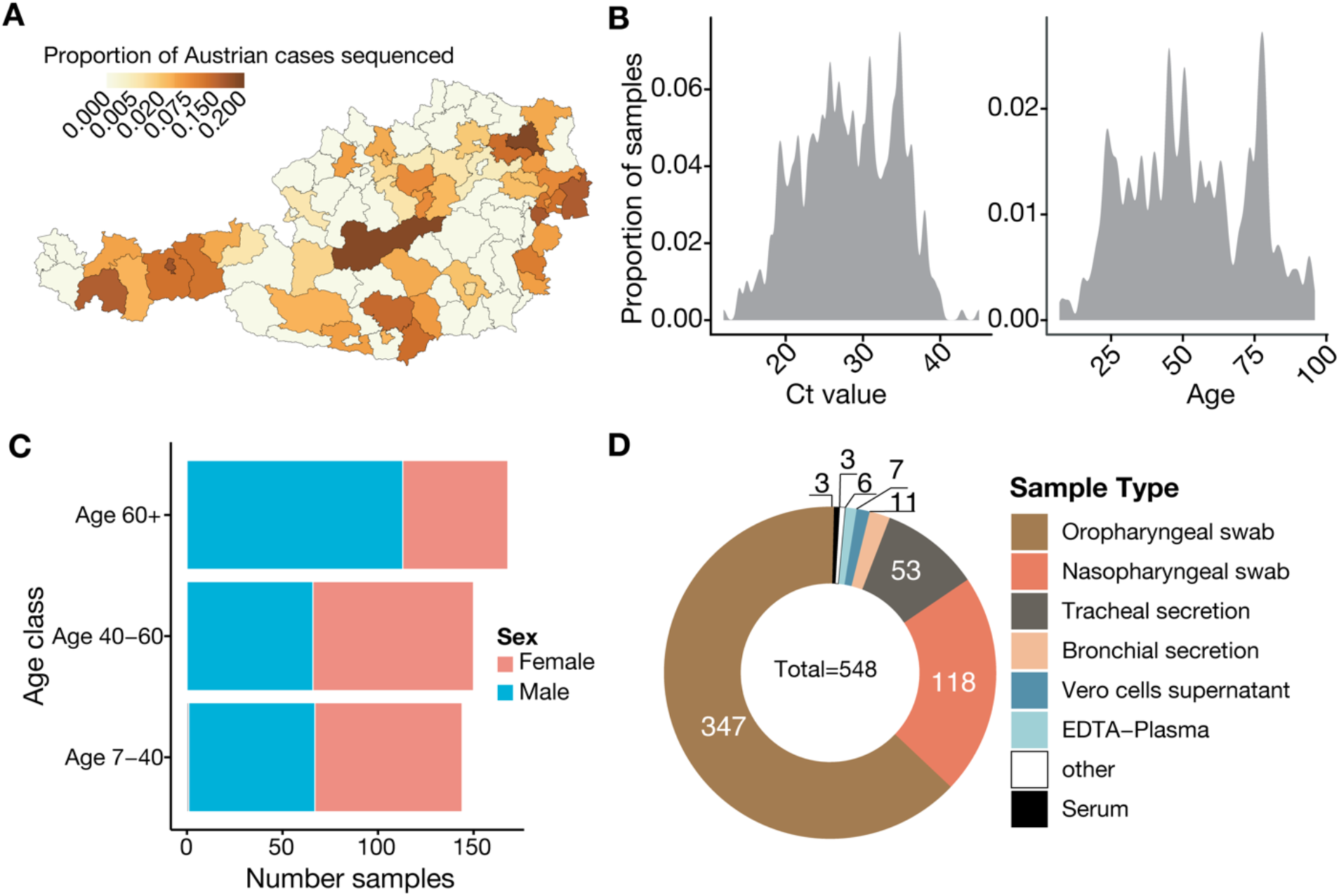
Data overview. (**A**) Proportion of sequenced SARS-CoV-2 samples among positive cases reported across Austrian districts. (**B**) Distribution of patients’ Ct values and age across the sequenced samples. (**C**) Number of female and male patient samples as a function of age class. (**D**) Number of samples for each sampling type category.

**Fig. S2.**
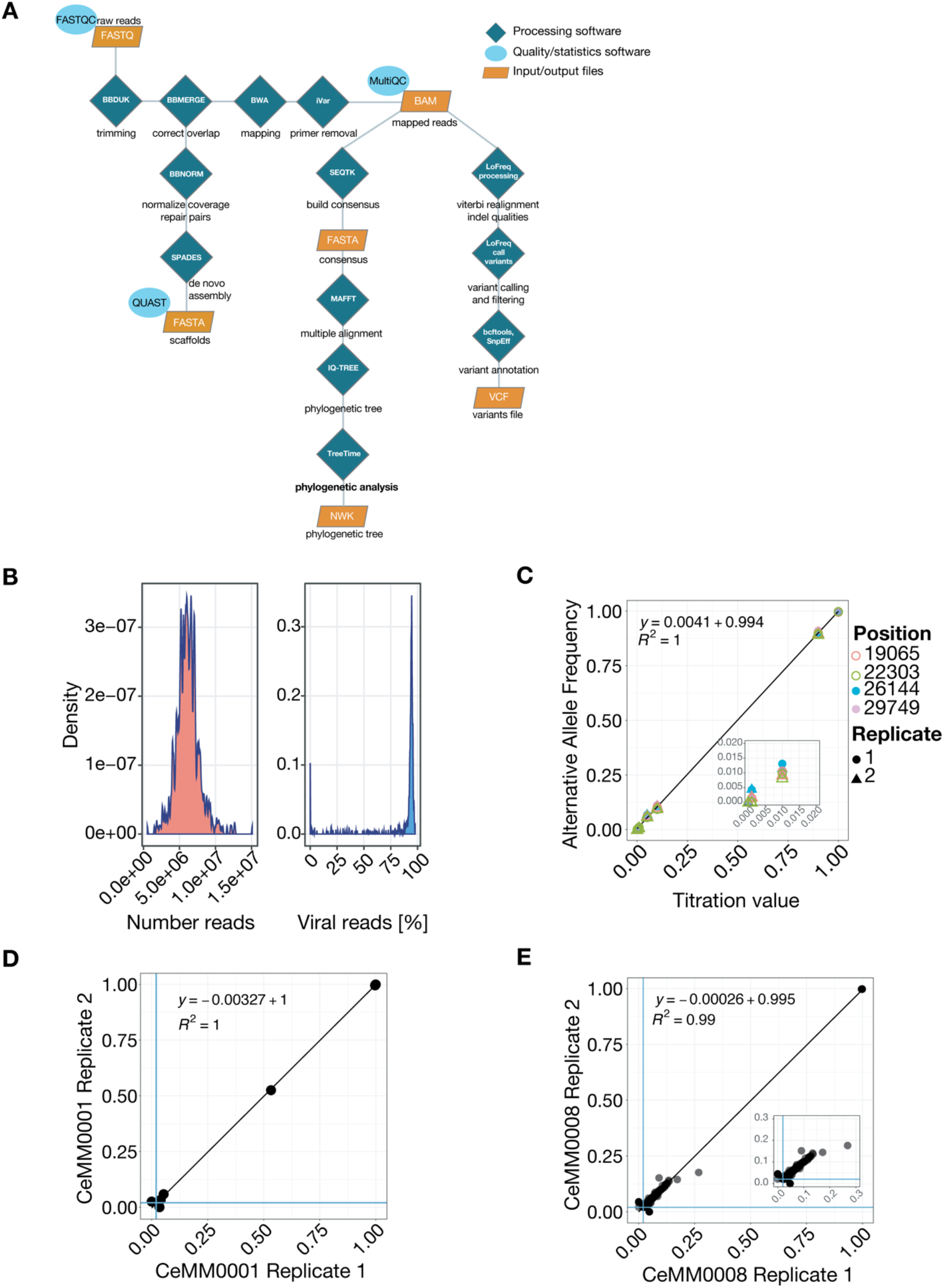
Technical pipeline and controls. (A) Processing pipeline from raw sequencing reads to fasta genomes, phylogenetic trees and low frequency mutation calling. (B) Distribution of the number of reads and the percentage of viral reads for all sequenced samples. (**C**) Mixture of two synthetic viral genomes in increasing ratios (0.1%, 1%, 5%, 10%, 90% and 100%). The two technical replicates of this titration are depicted with different symbols. (**D**) Comparison of variant detection for two independent full processing (PCR amplification, library preparation, sequencing) of the same patient sample, CeMM0001. (**E**) Comparison of variant detection for two independent sequencing runs of the same patient sample CeMM0008.

**Fig. S3.**
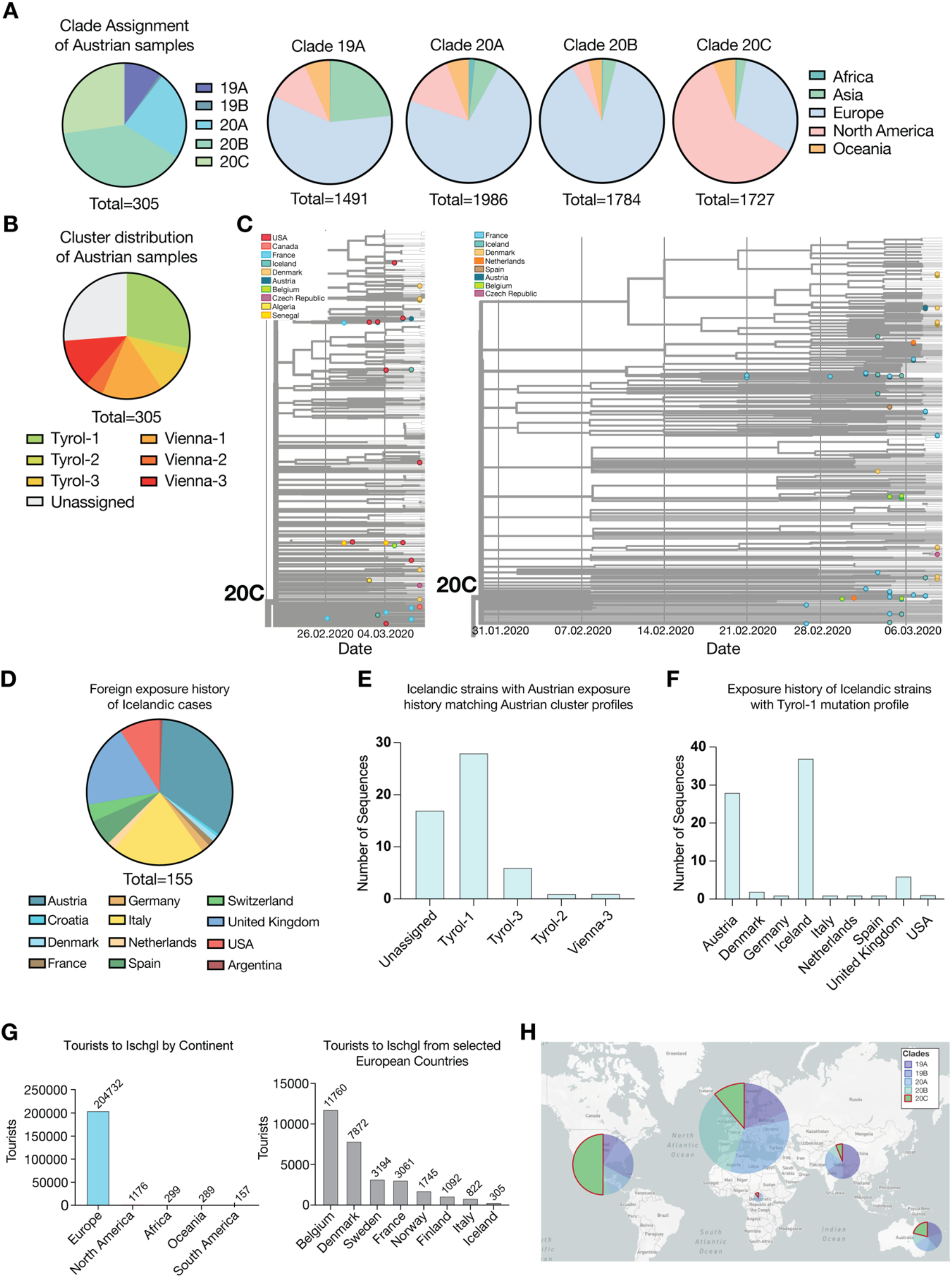
Phylogenetic analysis of SARS-CoV-2 sequences from Austrian COVID-19 patients in global context. (**A**) Nextstrain clade assignment of Austrian samples and geographic distribution of strains in the five clades defined by Nextstrain. The analysis of the geographic distribution of clades bases on information for 8,000 strains in the global phylogenetic analysis in this study. (**B**) Distribution of SARS-CoV-2 from Austrian COVID-19 sequences over the six phylogenetic clusters identified in this publication. (**C**) Clade 20C of time-resolved phylogenetic trees reconstructed from 7,695 randomly subsampled global strains and 305 Austrian strains (left) or all 7675 European high-quality sequences dated before 31^st^ of March. (**D**) Statistics of foreign exposure history of Icelandic COVID-19 cases as reported in GISAID. (**E**) Icelandic strains with Austrian exposure history matching Austrian cluster profiles. (**F**) Exposure history of all SARS-CoV-2 sequences from Icelandic COVID-19 cases available on GISAID that match the mutation profile of the phylogenetic cluster Tyrol-1. (**G**) International tourists visiting Ischgl between December 2019 and March 2020 by continent and selected European countries. (**H**) Distribution of SARS-CoV-2 strains over global clades across continents.

**Fig. S4.**
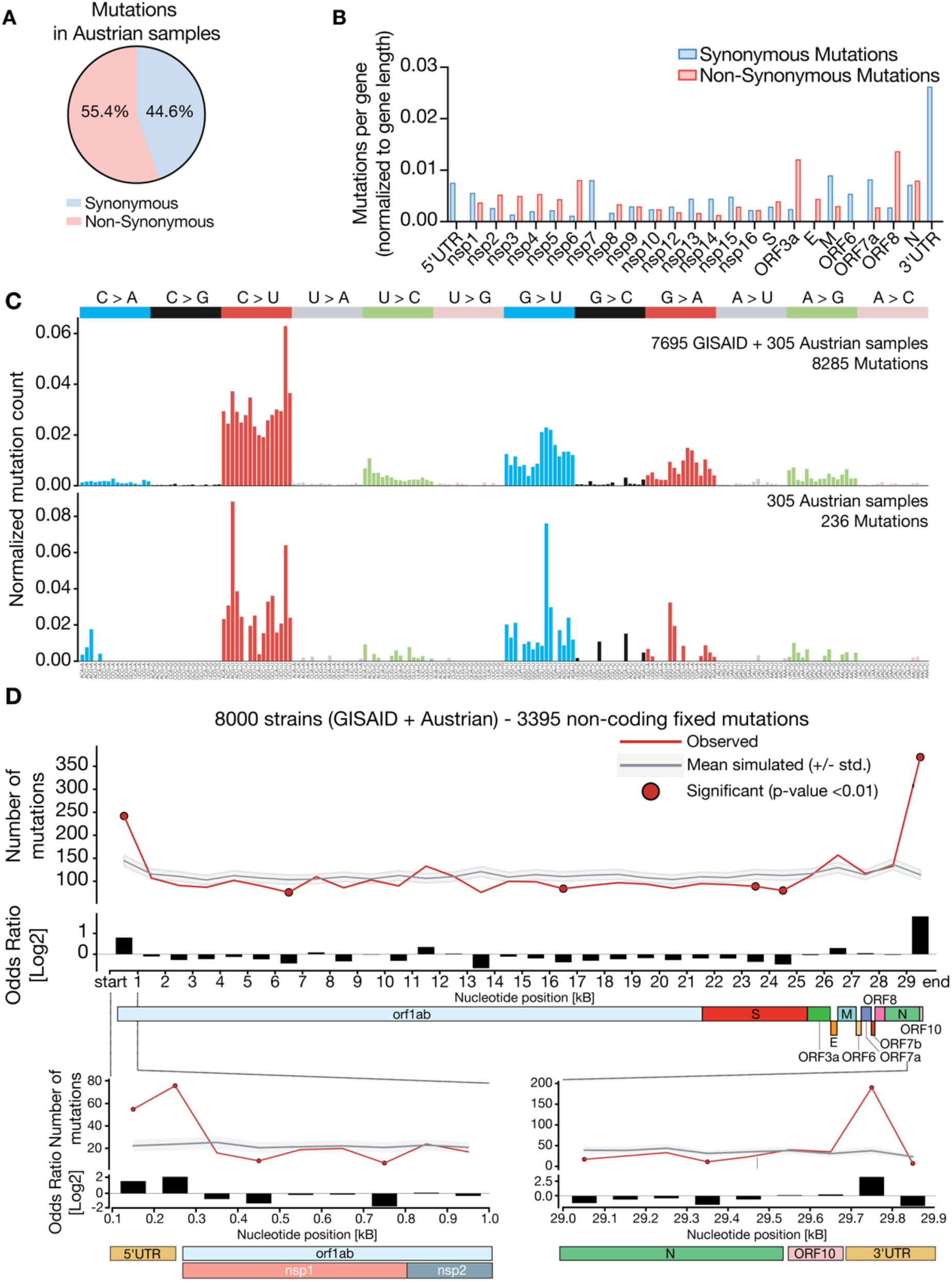
Mutational analysis of fixed mutations in SARS-CoV-2 sequences. (**A**) Ratio of non-synonymous to synonymous mutations in unique mutations identified in Austrian SARS-CoV-2 sequences. (**B**) Frequencies of synonymous and non-synonymous mutations per gene or genomic region normalized to length of the respective gene, genomic region or gene product (nsp1-16). (**C**) Mutational spectra panel. Mutational profile of interhost mutations. Relative probability of each trinucleotide change for mutations across SARS-CoV-2 sequences in 7,695 global sequences obtained from GISAID samples plus 305 Austrian samples (top) or 305 SARS-CoV-2 sequences from Austrian COVID-19 patients (bottom). (**D**) Mutation rate distribution along the SARS-CoV-2 genome. Top panel shows a 1kb window comparison of the observed number of synonymous mutations across the global subsample of 7,695 SARS-CoV-2 sequences from GISAID compared to the expected distribution (based on 10^6^ randomizations) according to their tri-nucleotide context. The grey line indicates the mean number of simulated mutations in the window, the colored background represents the distribution of expected mutations (+/- standard deviation with respect to the mean) and the red dots indicate a significant difference (G-test goodness of fit p-value <0.01). Odds ratio in log2 scale of the observed compared to expected number of synonymous mutations across the thirty 1kb windows of the SARS-CoV-2 genome. The bottom panel shows a zoom-in into the mutation rate across the first (left) and last (right) 1kb windows. The comparisons were performed using ten 100bp windows. Gene annotations for SARS-CoV-2 genome are given below the top panel.

**Fig. S5:**
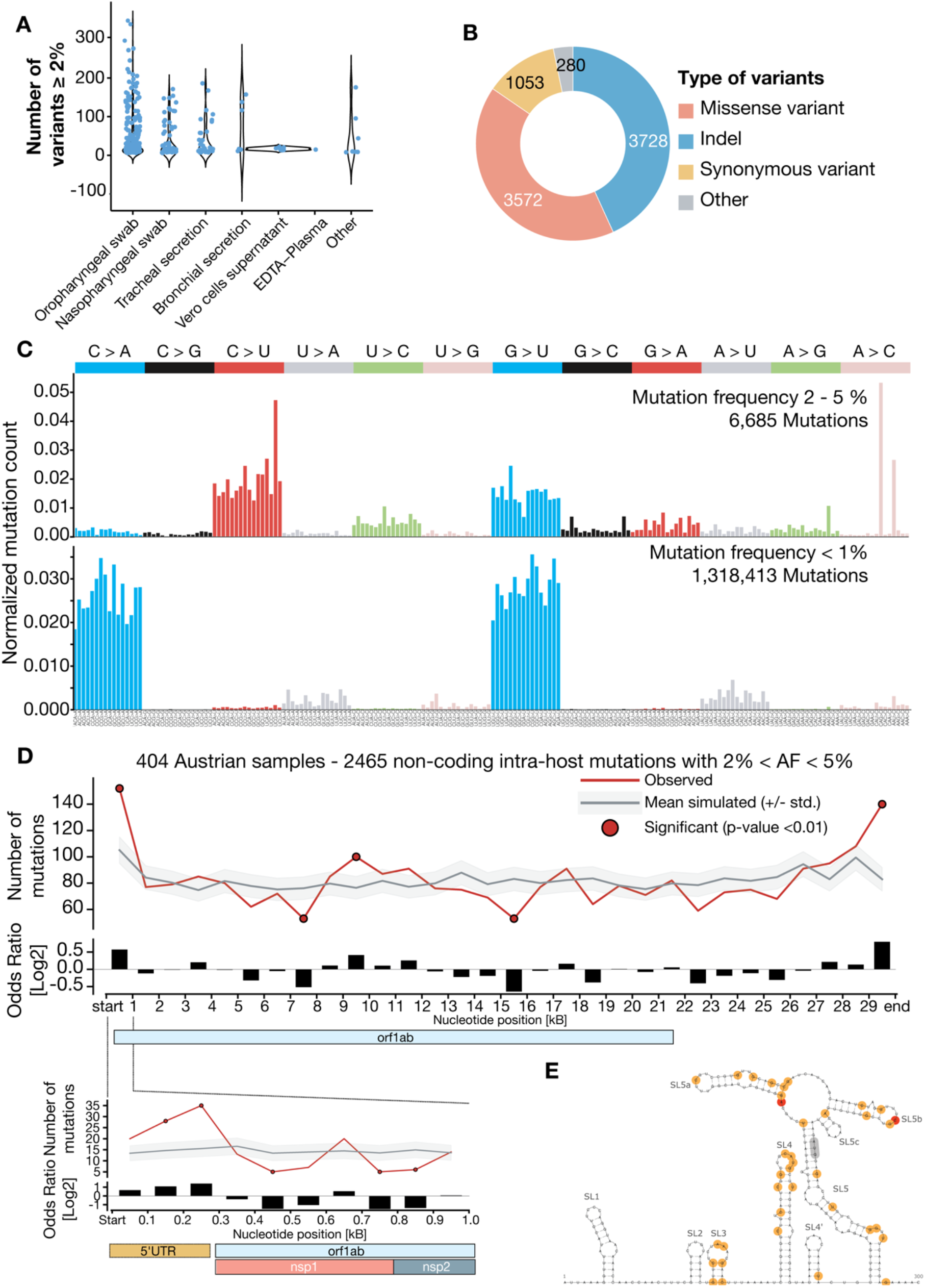
Analysis of low-frequency mutations. (**A**) Number of variants detected across the different sample types. (**B**) Number of variants per variant class. (**C**) Mutational profile (relative probability of each trinucleotide) of 6,685 intra-host mutations across Austrian samples (Alleles Frequencies between 0.02 and 0.05) (upper panel). Mutational profile (relative probability of each trinucleotide) of 1,318,413 intra-host mutations across Austrian samples (Alleles Frequencies less than 0.01) (lower panel). (**D**) Analysis of the mutation rate (analogous to the interhost mutation rate panel) across the SARS-CoV-2 genome using 2465 intra-host non-protein affecting mutations with alleles frequencies between 0.02 and 0.5. (**E**) RNA secondary structure prediction of the upstream 300nt of the SARS-CoV-2 reference genome (NC 045512.2), comprising the complete 5’untranslated region (UTR) and parts of the nsp1 protein nucleotide sequence. The canonical AUG start codon is located in a stacked region of SL5 (highlighted in gray). Mutational hotspots observed in the Austrian SARS-CoV-2 samples are highlighted in color. Two fixed mutations at positions 187 and 241, respectively, are marked in red. Low-frequency variants with an abundance between 2% and 50% in individual samples are shown in orange. Insertion and deletion variants are not shown.

**Fig. S6:**
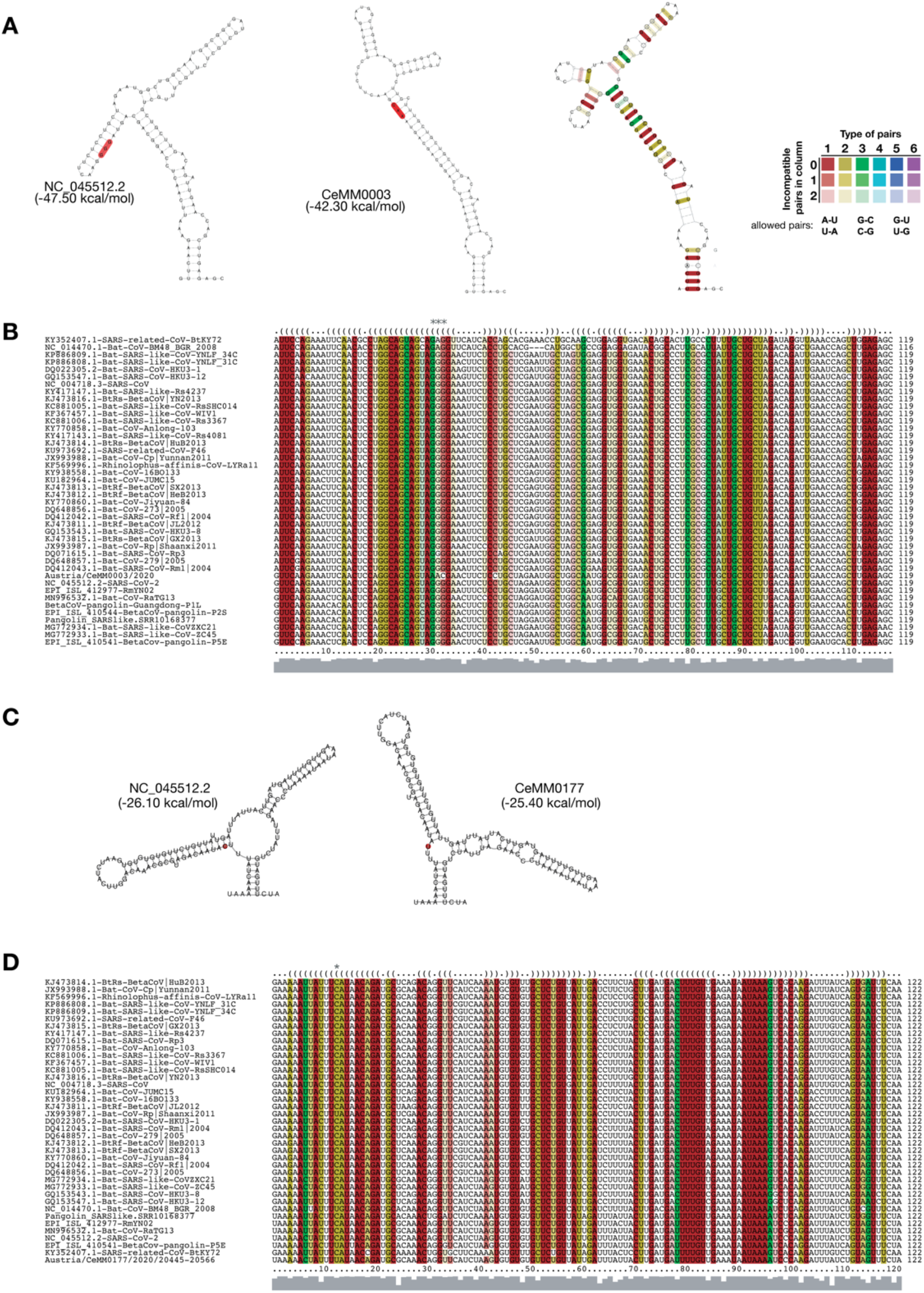
Predictions of mutational impact on RNA secondary structure. (**A**) Impact of a triple nucleotide mutation at positions 28881-3 on the RNA secondary structure. Minimum free energy (MFE) structure predictions of the 119nt region are shown for the SARS-CoV-2 reference genome NC 045512.2 as well as the mutation in sample CeMM0003. Consensus structure prediction is also reported. Thermodynamic stability of each structure is reported between brackets. (**B**) Corresponding structural multiple sequence alignment of the 119nt around the triple mutation at positions 28881-3 window in related Sarbecovirus genomes. The mutation site is marked by three asterisks above the dot-bracket consensus structure in the alignment. Colors following RNAalifold coloring scheme, indicating different levels of covariation for individual base pairs. (**C**) Impact of mutation at position 20,457 on the RNA secondary structure. MFE structure prediction of the 122nt region in the SARS-CoV-2 reference genome NC 045512.2 with the mutation site at position 13 highlighted in red (left). MFE prediction of the C>U variant as observed in sample CeMM0177 (right). Thermodynamic stability of each structure is reported between brackets. (**D**) Corresponding structural multiple sequence alignment of the 122nt window around the mutation at position 20457 in related Sarbecovirus genomes. The mutation site is marked by one asterisk above the dot-bracket consensus structure in the alignment. Colors following RNAalifold coloring scheme, indicating different levels of covariation for individual base pairs.

**Fig. S7:**
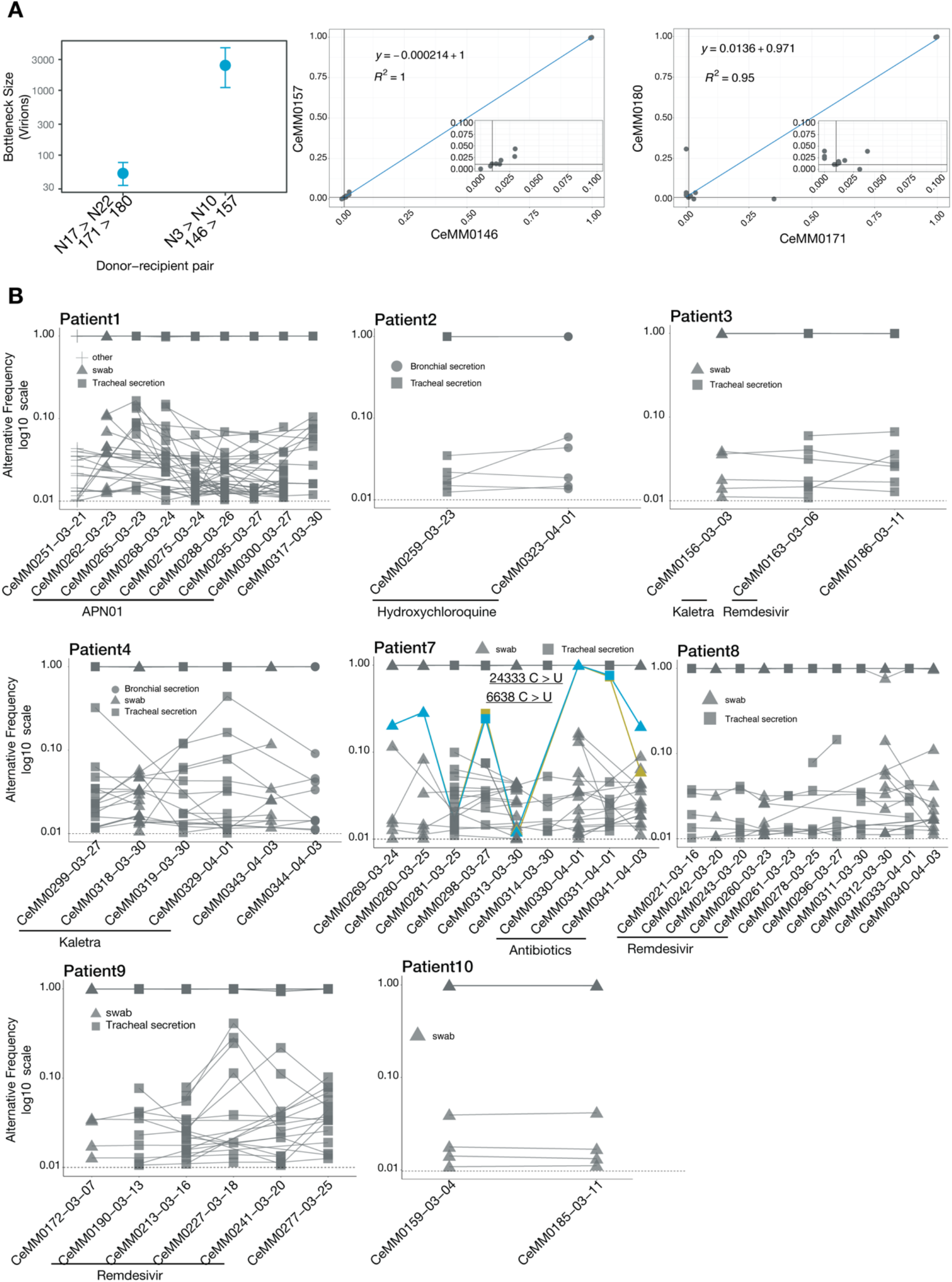
Viral intra-host diversity in individual patients. (**A**) Two examples of low (15 variants) and high bottleneck (14 variants) estimation in cluster A between N17 > N22. Samples 171 > 180 for low bottleneck and 146 > 157 for high bottleneck estimate. (**B**) Alternative allele frequency of all variants across time points related to patients 1, 2, 3, 4, 7, 8, 9, and 10. Only variants shared at between at least two time points are represented.

## Supplementary Tables

**Table S1: Acknowledgements for SARS-CoV-2 genome sequences derived from GISAID**

**Table S2: Epidemiological clusters referred to in this study.**

**Table S3: Clinical information of COVID-19 patients relating to Figure 3 and Figure S7.**

## Notes

https://github.com/Bergthalerlab/SARSCoV2_Code

### Competing Interest Statement

The authors have declared no competing interest.

